# Fitness traits of deoxynivalenol and nivalenol-producing Fusarium graminearum species complex strains from wheat

**DOI:** 10.1101/235895

**Authors:** C.P. Nicolli, F.J. Machado, P. Spolti, E.M. Del Ponte

**Affiliations:** Departamento de Fitopatologia, Universidade Federal de Vicosa, 36570-000, Vicosa, MG, Brazil.

**Keywords:** *Triticum aestivum*, trichothecenes, fitness, FGSC

## Abstract

*Fusarium graminearum* of the 15-acetyl(A)deoxynivalenol(D0N) chemotype is the main cause of Fusarium head blight (FHB) of wheat in south of Brazil. However, 3-ADON and nivalenol(NIV) chemotypes have been found in other members of the species complex causing FHB in wheat. To improve our understanding of the pathogen ecology, we assessed a range of fitness-related traits in a sample of 30 strains representatives of 15-ADON (*F. graminearum*), 3-ADON (*F. cortaderiae* and *F. austroamericanum*) and NIV (*F. meridionale* and F. *cortaderiae*). These included: perithecia formation on three cereal-based substrates, mycelial growth at two suboptimal temperatures, sporulation and germination, pathogenicity towards a susceptible and a moderately resistant cultivar and sensitivity to tebuconazole. The most important trait favoring *F. graminearum* was its 2x higher sexual fertility (> 40% PPI = perithecia production index) than the other species (< 30% PPI); PPI varied among substrates (maize > rice > wheat). In addition, sensitivity to tebuconazole appeared lower in *F. graminearum* which had the only strain with EC50 > 1 ppm. In the pathogenicity assays, the DON-producers were generally more aggressive (1.5 to 2x higher final severity) towards the two cultivars, with 3-ADON or 15-ADON leading to higher area under the severity curve than the NIV strains in the susceptible and moderately resistant cv., respectively. There was significant variation among strains of a same species with regards asexual fertility (mycelial growth, macroconidia production and germination), which suggest a strain-rather than a species-specific differences. These results contribute new knowledge to improve our understanding of the pathogen-related traits that may explain the dominance of certain members of the species complex in specific wheat agroecosystems.

## Introduction

Fusarium head blight (FHB) is a major disease of wheat and small grain crops due to the reduction in yield and contamination of grain with dangerous mycotoxins (McMullen et al. 2012). Deoxynivalenol (DON) is the most significant mycotoxin given its widespread occurrence at levels of concern to human and animal health (Tanaka et al. 1988). However, nivalenol (NIV), another type-B trichothecene, has been found in commercial wheat samples from Brazil (Del Ponte et al. 2015) and also in inoculations of wheat under controlled conditions (Mendes et al. 2018; Nicolli et al. 2015). A molecular survey conducted during the last decade showed that *Fusarium graminearum* (O’Donnell et al. 2004), formerly lineage 7 (O’Donnell et al. 2000), is the main (>85% frequency) *F. graminearum* species complex (FGSC) member causing FHB of wheat in Brazil (Del Ponte et al. 2015). The *F. graminearum* strains found in Brazilian wheat and barley analyzed so far are exclusively of the 15-acetyl(A)DON genotype, a mainly DON producer (Astolfi et al. 2012; Castañares et al. 2016; Del Ponte et al. 2015). *Fusarium meridionale* is the second most dominant FHB pathogen (>15% overall) and an important regional contributor of FHB in the major wheat-producing region of Brazil, Paraná State, where its frequency increases to ~30% in wheat spikes and grain (Del Ponte et al. 2015). This species is the main cause (> 96% frequency) of Gibberella ear rot and stalk rot as well the most frequent species surviving on maize stubbles, a large reservoir of inoculum for FHB epidemics (Del Ponte et al. 2015; Kühnem et al. 2015). Other FGSC members that occur at smaller frequency in wheat include *F. cortaderiae* and *F. austroamericanum* that possesses either 3ADON or NIV trichothecene genotype (Del Ponte et al. 2015). Nivalenol, as well as 3-ADON strains within FGSC members are actually much less common in wheat grown in the Americas, especially in Brazil (Del Ponte et al. 2015), but there are concerns about their ability to produce different or greater amount of trichothecenes (Mendes et al. 2018; Umpiérrez-Failache et al. 2013) and potential shifts in climate that may favor build up of inoculum of the currently minor species (Juroszek et al. 2013; Vaughan et al. 2016). In the state of Louisiana, U.S., NIV-type populations of *F. graminearum*, which were found at high frequency (79%) among isolates from small-grain-growing regions, accumulated four times less trichothecene toxin than DON types on inoculated wheat (Gale et al. 2011). The 3-ADON strains of *F. graminearum*, which appeared to emerge and spread into some regions of North America, were able to produce higher quantities of total trichothecenes and exhibit higher fecundity and growth rates than 15-ADON strains (Ward et al. 2008). More recently, the aggressiveness of *F. graminearum* strains of the 15ADON or 3-ADON genotypes from New York towards a susceptible wheat cultivar did not differ, thus suggesting no apparent advantage of 3-ADON compared to 15-ADON genotype, although the former produced significantly greater amount of trichothecenes (Spolti et al. 2014a).

In the south of Brazil, besides the dominant *F. graminearum* of the 15-ADON chemotype population, 3-ADON strains were found only in *F. cortaderiae* and *F. austroamericanum* isolates, while the NIV genotype was found in *F. cortaderiae*, *F. meridionale* and *F. asiaticum* (Del Ponte et al. 2015). The 3-ADON strains from wheat were found at higher elevation regions with cooler winter and spring seasons, which suggested that climate could also shape species composition, besides cereal hosts (Del Ponte et al. 2015). A recent study comparing 15-ADON and NIV strains of *F. graminearum* from China showed that the former grew faster at both 20 oC and 25 oC, produced higher amount of perithecia, and were more aggressive towards a wheat cultivar (Liu et al. 2017). These findings of enhanced aggressiveness of the 15-ADON agree with studies in Brazil, although the NIV strains were of a different species (*F. meridionale*) (Spolti et al. 2012a; Spolti et al. 2013). The toxigenic potential of *F. graminearum* also appeared to be higher than F. meridionale because much greater amount of DON was produced compared to NIV produced by the latter (Nicolli et al. 2015).

Empirical knowledge of some components of the life cycle of other FGSC members than *F. graminearum* and *F. meridionale* in Brazil, such as the saprophytic ability, and whether this is influenced by the cereal-based substrate and temperature, which may provide competitive advantages for a species to build up inoculum for further spread are not available. In this study, a sample of 30 strains representative of three trichothecene genotypes among four FGSC members associated with FHB of wheat in southern Brazil was selected to investigate the fitness-related traits related to both the saprophytic and pathogenic phase.

## Material and Methods

### Isolates and experiments

The thirty isolates selected for this study were originated from surveys of symptomatic wheat spikes in commercial fields during four seasons (2007/08 to 2011/12) in southern Brazil. These were accurately identified to species and trichothecenes genotypes using PCR-based and multilocus-genotyping (MLGT) assays (Astolfi et al. 2012; Del Ponte et al. 2015). The strains were deposited and are available upon request at a mycological collection (CML - “Coleção Micológica de Lavras”). Table 1 shows detailed information for these strains and indicates in which experiments they were used. A total of five different experiments were conducted to assess saprophytic (sexual and asexual reproduction), pathogenic (severity towards two cultivars), and sensitivity to a triazole fungicide (tebuconazole).

**Table 1.** Information for a collection of 30 strains belonging to four phylogenetic species of the *Fusarium graminearum* species complex isolated from symptomatic wheat heads from fields in Rio Grande do Sul State, Brazil, which were used in five different experiments in this study

| CML a | Location | Year | Species | Genotypeb | Experimentc |
| --- | --- | --- | --- | --- | --- |
| 3065 | Panambi | 2009 | *F. graminearum* | 15-ADON | 2, 3, 4 |
| 3066 | Ijuí | 2009 | *F. graminearum* | 15-ADON | 1,4 |
| 3067 | Carazinho | 2010 | *F. graminearum* | 15-ADON | 1,2,3,4 |
| 3071 | Tapejara | 2010 | *F. graminearum* | 15-ADON | 1,2,3,4,5 |
| 3069 | Coxilha | 2010 | *F. graminearum* | 15-ADON | 1, 2, 4,5 |
| 3070 | Santa Bárbara do Sul | 2011 | *F. graminearum* | 15-ADON | 1, 2, 4 |
| 3830 | Não-Me-Toque | 2011 | *F. graminearum* | 15-ADON | 1,2,3 |
| 3833 | Não-Me-Toque | 2011 | *F. graminearum* | 15-ADON | 1 |
| 3064 | Cruz Alta | 2007 | *F. graminearum* | 15-ADON | 1,4,5 |
| 3068 | Ernestina | 2007 | *F. graminearum* | 15-ADON | 1,2,3,4,5 |
| 3382 | Caseiros | 2010 | *F. austroamericanum* | 3-ADON | 1,2,3,5 |
| 3384 | Barreto | 2010 | *F. austroamericanum* | 3-ADON | 1,2,3,4,5 |
| 3379 | Carazinho | 2010 | *F. cortaderiae* | 3-ADON | 1,2,3,4,5 |
| 3845 | Lagoa Vermelha | 2010 | *F. cortaderiae* | 3-ADON | 1,2,3,4 |
| 3841 | Lagoa Vermelha | 2010 | *F. cortaderiae* | 3-ADON | 1,2,3,4,5 |
| 3831 | Panambi | 2009 | *F. cortaderiae* | NIV | 1,2,3,4,5 |
| 3836 | Coronel Barros | 2009 | *F. cortaderiae* | NIV | 1,2,4 |
| 3837 | Coronel Barros | 2009 | *F. cortaderiae* | NIV | 1,2,3,4,5 |
| 3840 | Mato Castelhano | 2010 | *F. cortaderiae* | NIV | 1,2,4 |
| 3834 | Barreto | 2010 | *F. cortaderiae* | NIV | 1,2,4 |
| 3378 | Vacaria | 2010 | *F. cortaderiae* | NIV | 1,2,3,4,5 |
| 3835 | Ernestina | 2007 | *F. cortaderiae* | NIV | 1,2,3,4 |
| 3375 | Santa Bárbara do Sul | 2009 | *F. meridionale* | NIV | 1,2,4 |
| 3843 | Ernestina | 2009 | *F. meridionale* | NIV | 1,2,3,4 |
| 3344 | Coxilha | 2009 | *F. meridionale* | NIV | 1,2,3,4,5 |
| 3381 | Caseiros | 2010 | *F. meridionale* | NIV | 1,2,3,4,5 |
| 3839 | Marau | 2011 | *F. meridionale* | NIV | 1,2,3,4,5 |
| 3377 | Sertão | 2011 | *F. meridionale* | NIV | 1,2,3,4 |
| 3838 | Nonoai | 2007 | *F. meridionale* | NIV | 1,2,3,4 |
| 3374 | Nonoai | 2007 | *F. meridionale* | NIV | 1,2,4 |

a Culture collection code: CML = Coleção Micológica de Lavras, Universidade Federal de Lavras, Lavras, Minas Gerais, Brazil. b Trichothecene genotype identified by the Luminex method: 15-ADON = 15-acetyl-deoxynivalenol; 3-ADON = 3-acetyl-deoxynivalenol; NIV =
nivalenol. c Experiments: 1 = Perithecial production; 2 = Mycelial growth; 3 = Sporulation and germination; 4 = Pathogenicity; 5 = Sensitivity to tebuconazole.

### Perithecial production

Perithecial production, as indicative of sexual fertility, was evaluated using kernel-based substrates composed of a distinct cereal host: wheat, rice or maize. The method was adapted from Chen & Zhou (2009). Briefly, 20g of grain was soaked in 10 mL distilled water and autoclaved for 20 min at 127 o C daily during consecutive days. Afterwards, kernels were inoculated with three discs (6 mm diameter) of mycelia and kept in the dark for 21 days at 25 oC. Then, the colonized grains were transferred to a bench in a greenhouse with controlled temperature (25 ±3 oC) where they remained for seven days for drying. After drying, they were placed in plastic box (11 × 11 × 3.5 cm) containing sterile sand layer (2 cm) and three sheets of filter paper on top of the sand. The filter paper was moistened daily with sterile distilled water. After 21 days, 50 grains were randomly selected from each substrate-isolate combination and individually scored visually using a diagrammatic key developed for this study and which contains four scores representing a percent range for the grain surface covered with perithecia (Supplementary Fig. 1). The frequency of the scores for each species-genotype combination (five groups) was normalized to a perithecial production index (PPI, %).

### Mycelial growth

Isolates grown on PDA (20 mL per dish) for seven days at 25 oC in continuous darkness provided mycelial discs (6 mm diameter) which were individually placed in the center of plastic Petri dishes (90 mm diameter). The cultures were incubated in growth chambers at two regimes of sub-optimal constant temperature (15 oC or 30 oC) under continuous darkness for five days when the diameter of the colony was measured from two perpendicular directions. The mycelial growth rate (mm/day) was calculated after subtracting the mycelium plug size (Spolti et al. 2012b). Three replicates (plates) were used per treatment.

### Sporulation and germination

The asexual fecundity of the strains was evaluated based on the macroconidia production in culture medium. Isolates were grown on SNA media for seven days under a 12/12 h light/dark cycle at 25 o C (Leslie and Summerell, 2006). After this period, three discs of 6 mm in diameter were removed from the edges of the developing colonies and immersed in 5 mL of sterile distilled water containing 0.1 mL Tween 0.001% in a test tube and shaken for 20 s. The spore numbers were counted and the sporulation expressed as number of spores/mL. Spore germination was observed for a subsample of 20 spores (three replicated slides) after six hours of exposure to water and results were expressed as percent of germinating spores.

### Pathogenicity

The pathogenicity of the strains was evaluated using a susceptible (BRS 194) and a moderately resistant (BRS Guamirim) cultivar. Seeds were sown in plastic pots (3 L) containing carbonized rice husk substrate (Tropstrato HT vegetables) fertilized with macronutrients (NPK 4-14-8). After seedling emergence, they were thinned to obtain 10 plants per pot. Inoculations were performed during the milk stage of grain development of the main tillers of the plants using the central-spikelet method (Engle et al. 2003). The mid-position spikelet (sixth from top to bottom) was marked with a pen and 20 μι of a spore suspension (105 conidia/mL) was gently applied inside the floret with the aid of a micropipette (Spolti et al. 2012b). After inoculation, the plants were kept at constant temperature (25 oC) to induce infection. The proportion of diseased spikelets in the wheat head was evaluated every three to five days during three weeks after inoculation. The experiment was repeated once in time and the data combined for analysis. The area under the diseased progress curve (AUDPC) was calculated and used to evaluate the effect of the trichothecene genotypes.

### Sensitivity to tebuconazole

Tebuconazole sensitivity was quantified by measuring mycelial growth on PDA media (potato dextrose agar) amended with increasing fungicide concentrations (Becher et al. 2010; Spolti et al. 2014b). Stock solution (100 μg active ingredient [a.i.]/mL) was adjusted by dilution of commercial formulation (Folicur 200 EC, BAYER S.A., 20% a.i.) in distilled. Stock solution was added to cooled PDA to obtain the following tested concentrations of 0.0; 0.5; 1.0; 2.0, 4.0 and 8.0 μg a.i./mL. One mycelia agar disc (6 mm in diameter) obtained from the edge of a 7-day-old culture was placed at the central position of a Petri dish (90 mm) containing 15 mL of PDA amended with the fungicide at each concentration. After four days of incubation at 25oC in darkness, radial growth was measured in two perpendicular directions and the disc diameter was subtracted. For each repeat of the experiment, two replicates were used. The effective fungicide concentration to reduce 50% of mycelial growth (EC50) was determined using a linear regression model as described elsewhere (Becher et al. 2010; Spolti et al. 2014b).

### Statistical analyses

For the data from the experiments to evaluate saprophytic growth (perithecia formation, mycelial growth and sporulation-germination) and pathogenicity, a multilevel linear mixed model was used to test the effects of the interaction, whenever present (e.g. temperature x genotype or species or species-genotype combination), or the single effect of a factor on the response variable at 0.05 significance level. In the model, replication for the isolates and isolates within each species-genotype or trichothecene genotype, depending on the experiment, were treated as random effects. The lsmeans and respective confidence intervals were obtained (using emmeans package of R) using kenward rogers df approximation. A Tukey method was used for comparing the means (alpha = 0.05) whenever the effect was deemed significant. Finally, the experiment to obtain EC50 values was repeated once in time and the data were combined for analysis. The means and confidence intervals were calculated.

## Results

### Saprophytic fitness

The proportion of grains with scores >1 (or >10% kernel area covered with perithecia) was apparently higher for maize and rice kernels substrates compared with wheat and for the 15-ADON isolates compared with NIV and 3-ADON (Figure 1A). All interactions tested in the model (species, genotypes or species-genotype combined versus substrates) significantly affected (P <0.001) perithecial production index (PPI), which was highest (>50%) for 15-ADON *F. graminearum* strains inoculated on rice and maize, and lowest for inoculations on wheat kernels (Figure 1B). There was a general trend of decreasing PPI in the wheat kernel substrate compared with the rice and maize substrates, as well as in the 3-ADON strains compared with the 15-ADON strains. For the two NIV-producing species, PPI was higher in F. meridionale strains than in F. cortaderiae strains inoculated in rice and maize, but not in wheat. There was no difference (*P* > 0.05) in PPI between the two 3-ADON species, regardless of the substrate.

**Figure 1.**
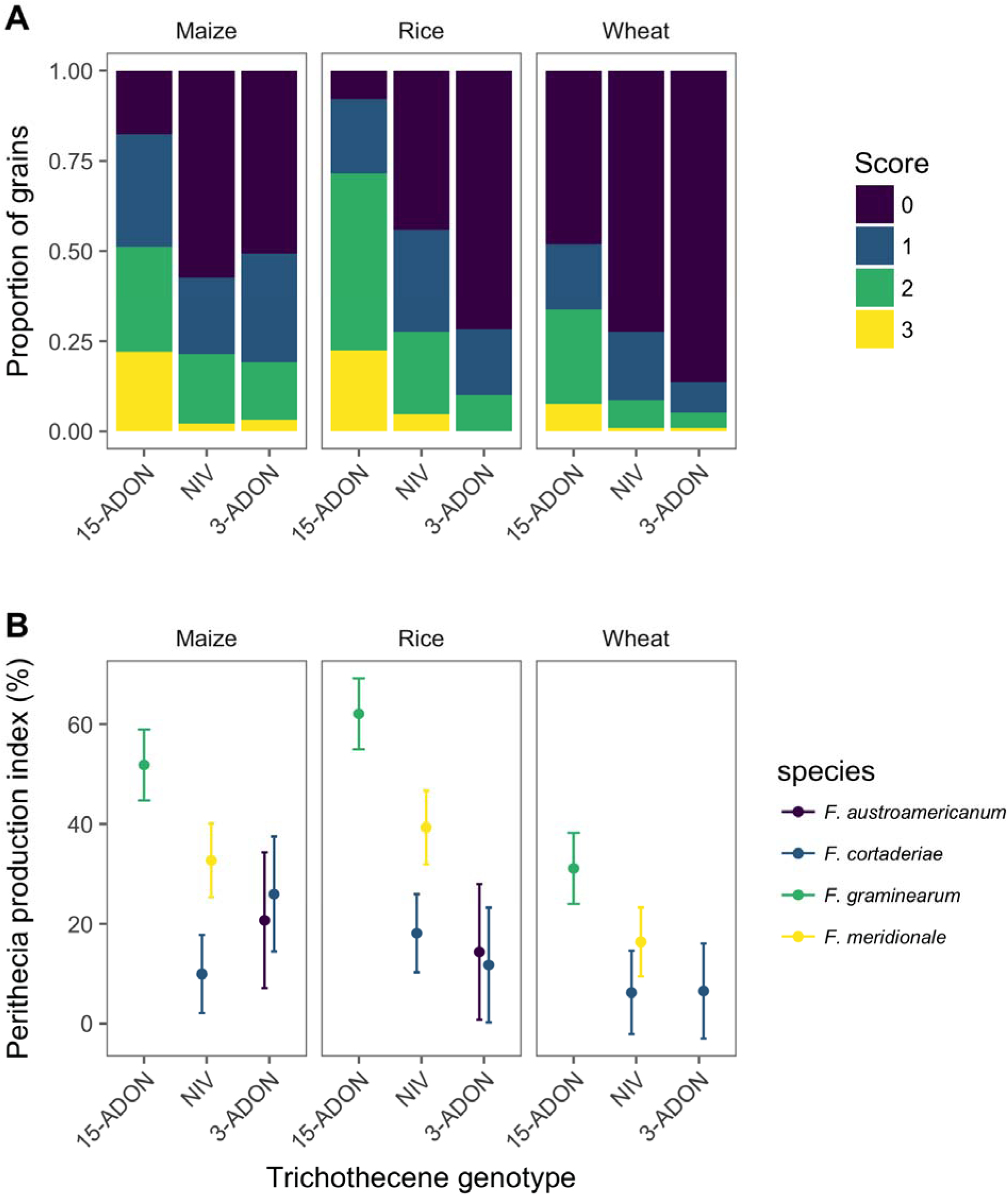
Relative frequency of perithecial production scores based on a visual ordinal (03) scale for the proportion of grains in each score for the three different cereal based substrates (wheat, rice or maize kernels) (A) and the least square means and 95% confidence interval for the perithecial production index which is a normalization of the frequency of the scores (B).

As expected, mean values of mycelial growth rate (MGR) was 84% higher at 30 oC (0.7 mm/day) than at 15 oC (1.2 mm/day) regardless of the trichothecene genotype. Within each of the temperatures, there were no differences in MGR means among the trichothecene genotypes (*P* > 0.20) as well as between species within trichothecene genotypes (Figure 2A). The variation in MGR was apparently higher within 15-ADON strains compared with the others, especially for the high temperature.

**Figure 2.**
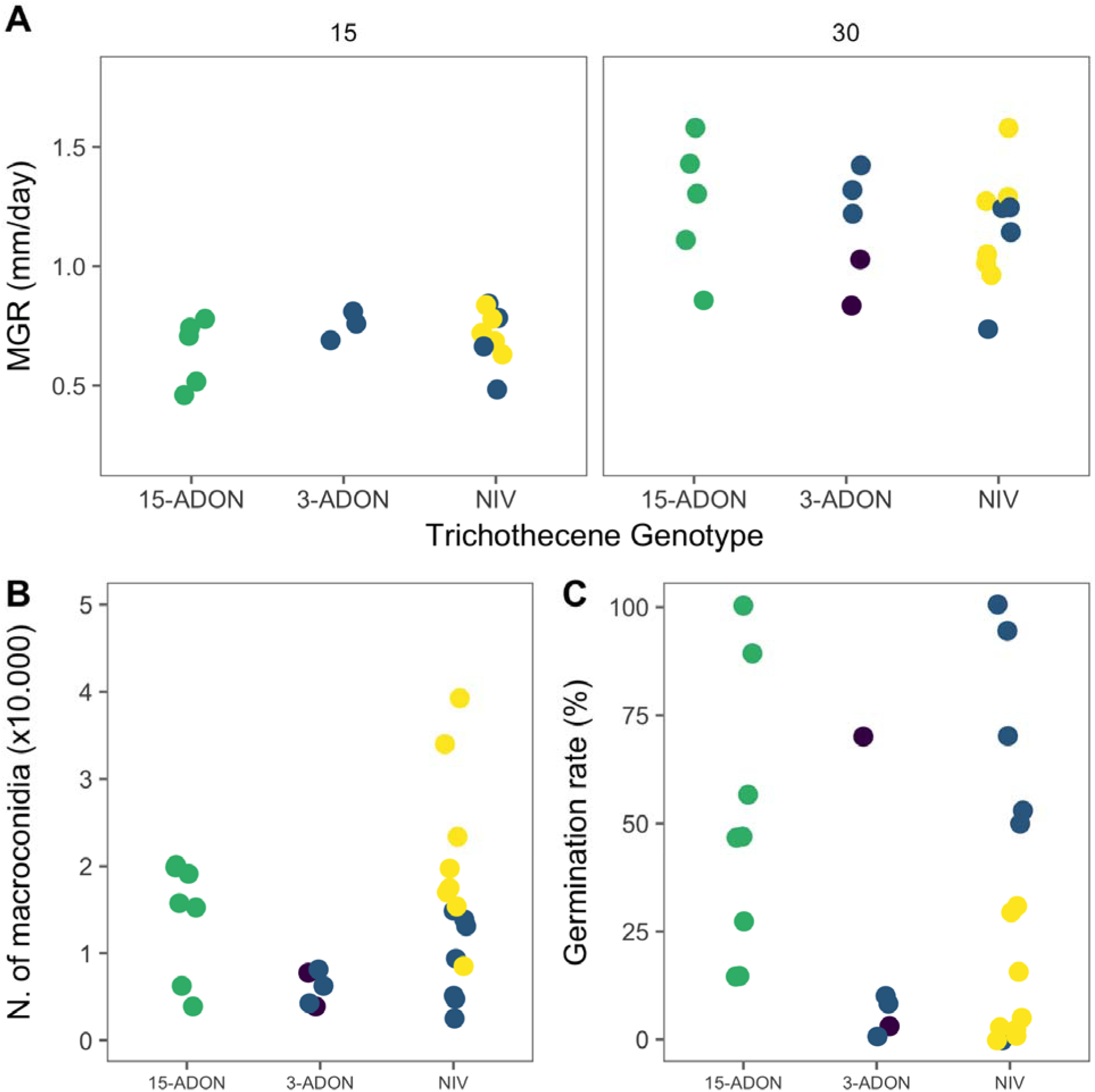
Mycelial growth and spore production by strains of four Fusarium graminearum species complex members possessing a nivalenol (NIV) or an acetylate form of deoxynivalenol (DON) genotype (3-ADON or 15-ADON) assessed based on mycelial growth at 15o C or 30o C (A), macroconidia production (B) and conidia germination (C).

All strains yielded macroconidia and there were differences in conidia production among the genotypes (*P* = 0.057), which were more evident when species-genotypes were compared (*P* < 0.001). The most prolific species was *F. meridionale* (2.18 × 103 spores/mL), which differed significantly from the other species (0.58 to 0.9 × 103 spores/mL), but not from *F. graminearum* (1.50 × 103 spores/mL) (Fig 2B).

The germination rate ranged from 0 to 100% across the strains. The greatest variability (80% range) was found for the *F. graminearum* strains, followed by F. austroamericanum for which the two strains differed by >70%. Therefore, the effect of genotype was not significant (P = 0.17), but when species-genotype was compared, the mean germination rate of F. meridionale (10.5%) differed significantly from *F. graminearum* (50%) and F. cortaderiae NIV (52.8%).

### Pathogenicity

FHB severity (proportion of symptomatic spikelets) increased over time and ranged from 10% to 40% in the two cultivars. Regardless of the species, disease progress appeared slower during the first ten days after inoculation (<10%) for the moderately resistant cultivar (BRS Guamirim) compared to the susceptible (BRS 194), for which the progress peaked and stabilized at 15% to 35% after 10 days (Figure 3). No formal test was used to compare these two cultivars because they were evaluated in different experiments.

**Figure 3.**
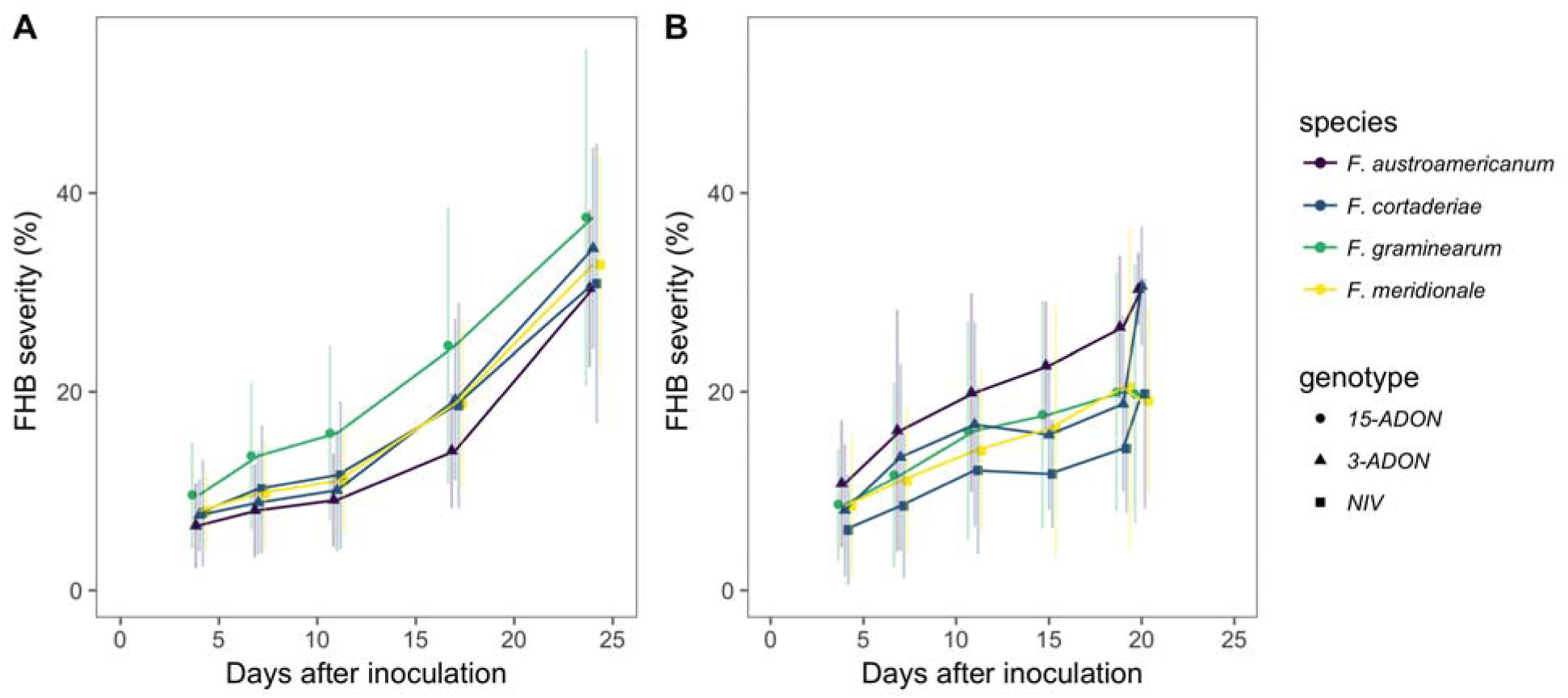
Mean (standard deviation) severity of Fusarium head blight (FHB) in a moderately resistant cultivar, BRS Guamirim (A), and a susceptible cultivar, BRS 194 (B), inoculated with strains of different Fusarium graminearum species complex members possessing a nivalenol (NIV) or an acetylate form of deoxynivalenol (DON) genotype (3-ADON or 15-ADON) found associated with FHB in Brazil.

Results of the mixed model to test the effect of genotype in the mean area under the severity curve showed the 15-ADON *F. graminearum* (432.1) was significantly higher (P < 0.05) than the 3-ADON-(331.03) and NIV-producing (319.6) species when inoculated in the moderately resistant cultivar, BRS Guamirim. On the other hand, the area under the severity curve in the BRS 194 inoculated with the 3-ADON genotypes (259.8, F. austroamericanum) was significantly higher than the NIV-producing strains (179.4); both not differing from the *F. graminearum* 15-ADON (204.38).

### Tebuconazole sensitivity

The estimated EC50 values ranged from 0.007 to 1.42 μg/mL across the strains. The highest mean EC50 value was found for one 15-ADON strain (1.42 μg/mL), while for the others the values were < 0.5 μg/mL (Fig. 4). Differences among the genotypes were not apparent across the strains and the limited number of samples prevented formal comparison.

**Figure 4.**
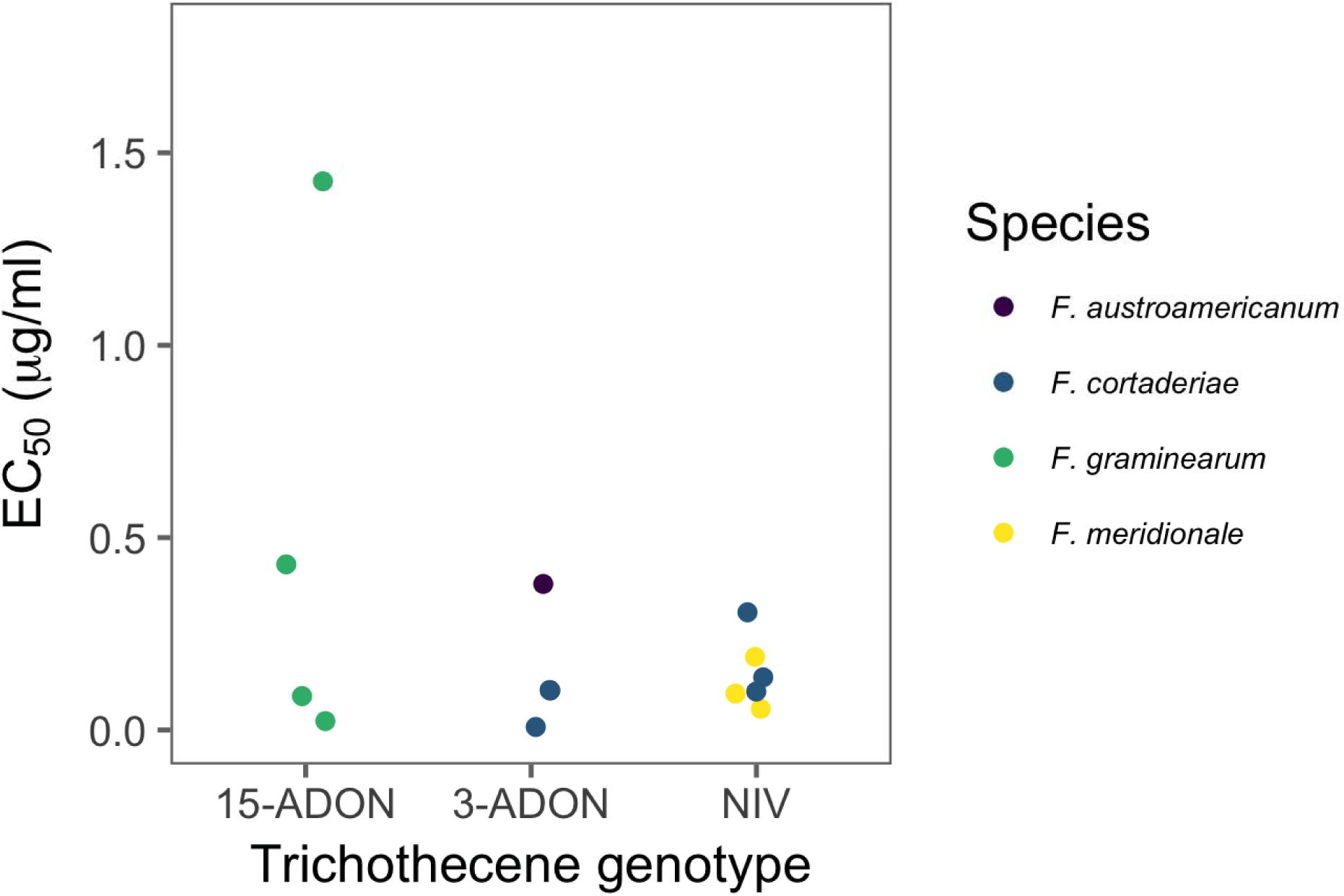
The effective concentration of tebuconazole that reduces 50% (EC50) of the mycelial growth of four Fusarium graminearum species complex members possessing a nivalenol (NIV) or an acetylate form of deoxynivalenol (DON) trichothecene genotype (3-ADON or 15-ADON) found associated with Fusarium head blight of wheat in Brazil.

## Discussion

In this study we investigated for the first time several traits related to different stages of the saprophytic and pathogenic phases of the FGSC life cycle using a sample of strains representative of four members (and three trichothecene chemotypes) associated with FHB in Brazil. At least for some components evaluated, results are supportive of the hypothesis that *F. graminearum* is more fit than the other species, which may help to explain its dominance as FHB pathogen of wheat in Brazil (Astolfi et al. 2012; Del Ponte et al. 2015; Scoz et al. 2009).

Our most striking result favoring *F. graminearum* was sexual fertility. These heterothallic fungi are able to overwinter in crop residues where they produce perithecia and ascospores which are able to disperse long-distance and infect wheat starting at flowering (Gilbert & Fernando, 2004; Khonga & Sutton, 2009; Pereyra & Dill-Macky, 2004). Thus, the perithecia production capacity should play an important role in the ecology and epidemiology (Leplat et al., 2013; McMullen et al, 2012). So far, few studies have characterized and compared FGSC members or chemotypes within a same species, from a same region/crop, with regards to perithecia and ascospore production (Lee et al. 2012; Liu et al. 2017; Spolti et al. 2014a). Similarly to others (Lee et al. 2009; Liu et al. 2017), we inferred on fertility based on the amount of perithecia produced on grain from three main cereal substrates. We found that *F. graminearum*, the only member that possess the 15-ADON trichothecene genotype in Brazil, produced higher amount of perithecia, followed by the NIV-producing *F. meridionale*, and then by the other two species with the 3-ADON or NIV chemotype. Interestingly, this ranking strongly agrees with that of the species frequency found in wheat and barley grain in the region (Astolfi et al., 2011; Astolfi et al., 2012; Del Ponte et al., 2015).

Previously, the ability of a dominant kernel-born FGSC member to produce significantly large amounts of perithecia in the original host was observed by Lee et al. (2009), who isolated 87% of *F. asiaticum* (main species found in kernels) strains from perithecia developing on rice plants after physiological maturity, followed by *F. graminearum* (13%). Recently, Liu et al (2017) compared perithecia production by *F. asiaticum* (NIV and 3-ADON) and *F. graminearum* (15-ADON) from China and found that the latter produced the highest amount of perithecia, which were visible two to three days earlier than the NIV and 3-ADON strains. In that Chinese region, *F. graminearum* 15-ADON is also dominant in wheat kernels. In New York, a comparison between 15-ADON and 3-ADON strains of *F. graminearum* from wheat showed no difference in saprophytic and pathogenic fitness towards wheat (Spolti et al. 2014a).

Previously, we found that perithecium-borne *F. meridionale* was highly prevalent in maize stubbles over the surface of two out of three wheat fields, and *F. graminearum* absent or at very low frequency in stubbles of all three fields (Del Ponte et al., 2015). We then hypothesized that the former was more fit towards maize, which was further confirmed more recently, as *F. meridionale* was largely dominant on stubbles, stalks and kernels of maize from Brazil (Kühnem et al. 2016). In the wheat study, although *F. meridionale* was dominant in perithecia from maize stubbles on the surface of wheat fields, it was the least frequent in wheat kernels from the same field, which were largely infected by *F. graminearum* (Del Ponte et al. 2015). Our results using three cereal-based substrates suggest that the dominance is not due to enhanced capacity of F. meridionale to produce perithecia when compared to *F. graminearum*, but to its ability to primarily cause stalk rot. Further studies are underway to investigate whether *F. meridionale* is able to outcompete *F. graminearum* in both single and mixed inoculations of maize stalks.

A few other questions remain. It would be instructive to further examine ascospore production and the aerobiology of these species in both controlled and field environment. In our previous field study, *F. meridionale* was the least frequent (<30%) species collected in selective media spore traps during wheat flowering (Del Ponte et al. 2015). Ideally, new studies should include more accurate or less-biased methods, such as molecular identification, given that during isolations, a fast-growing species may outcompete others. In addition, it would be important to monitor ascorporic inoculum over time and space when both species are present in similar frequency, similar to another study that used spore sampler placed near maize stalk residues bearing perithecia with mature ascospores (Manstretta & Rossi 2015). It is well know that FGSC are capable of long distance dispersal (Prussin et al. 2014) and that airborne populations above wheat canopies may be composed of well-mixed inoculum originated under the effect of turbulence and gravitational settling of inoculum from both local and external sources (Del Ponte et al. 2003; Schmale et al. 2006). The latter hypothesis has been suggested based on both aerobiological and spatial patterns studies in Brazil (Del Ponte et al. 2005; Spolti et al. 2015).

There was a large variation among the isolates from a same species-chemotype with regards to the response to the two sub-optimal temperatures tested in our study, thus preventing us from detecting differences in mycelial growth rate at the species or chemotype level. Previous studies have shown a large sensitivity of *F. graminearum* strains to temperature increase, which was indicative of potential for thermal adaptation (Backhouse et al. 2014; Spolti et al. 2014a; Zhan & McDonald, 2011) and saprophytic adaptability (Tunali et al. 2012). In our previous work evaluating mycelial growth rate of only two *F. graminearum* and two F. meridionale strains, growing at four increasing temperatures (10 to 30 oC), significant difference between species was found only at 25 oC and 30 oC, with *F. graminearum* growing faster than *F. meridionale* (Spolti et al. 2012a). Using a sample as low as eight strains, the maximum difference among strains of *F. graminearum* and *F. meridionale* found in this study was ~10 mm/day, which suggests caution when inferring species differences based on small sample.

Differences in macroconidia production in culture among species may be suggestive of advantages for further spread by assexual means (Leplat et al. 2013). We found that the variation in this trait was very large among isolates of the same species, especially *F. graminearum* and *F. meridionale*, the most represented species, which prevented us from detecting differences as found in our previous studies using a different set of isolates and smaller sample (Spolti et al. 2012a; Spolti & Del Ponte 2013). Recently, Liu et al. (2017) reported significantly greater macroconidia production by 20 *F. graminearum* 15-ADON strains than 20 *F. asiaticum* NIV strains, but they both did not differ from the F. asiaticum 3-ADON strains. The germination rate also varied considerably among strains of the same species, which makes largely unclear whether macroconidia production is a reliable variable to compare the asexual reproduction fitness among members of the complex given the inconsistencies and high variability. Moreover, in natural epidemics caused by FGSC, the importance of macroconidia inoculum is not clear, although studies highlighted the potential to dispersal and further infection (Paul et al. 2007).

We found that the species with the DON chemotype were generally more aggressive (higher area under severity curve) than the NIV ones, which is in agreement with our previous studies (Mendes et al. 2018; Spolti et al. 2012a). We could not compare statistically the effect of cultivar because the experiments were conducted at different times, and with different total time duration of the evaluations. Nevertheless, the area under the curve was in general higher for BRS 194, the susceptible one, compared with BRS Guamirim as shown earlier (Mendes et al. 2018; Spolti et al. 2012a). For the first time, we compared the two acetylates of DON within the Brazilian strains; while 15-ADON (*F. graminearum*) was the most aggressive in a susceptible cultivar, the 3-ADON (*F. austroamericanum*) was the most aggressive in the moderately resistant cultivar. Previously, studies comparing strains of the these two acetylates of DON showed differences in the rate of colonization, with 15-ADON being more aggressive than NIV, only in the less susceptible (BRS Guamirim), and not in BRS 194 (Spolti et al. 2012a).

We used the area under the curve of severity and not the final severity (last assessment) (Spolti et al. 2012a), or count of diseased spikelets at individual times (Liu et al. 2017) to compare the strains. Although these measures may be correlated, the area under the curve is more appropriate to capture differences due to variations in the daily progress (e.g. different curve shapes may lead to a same final severity). It would be important to standardize the methodology for assessing FHB pathogen aggressiveness, especially the type of variables and analysis when using central spikelet inoculation in order to compare data from multiple studies. These would include: inoculation timing, spore concentration, spikelet position, disease assessment, area under curve of final severity or disease count, etc. A previous study has shown that, in some cases, the bleaching above the inoculation point for some cultivars not due to fungal spread but to death of vascular tissue, which is hard to discern visually (Brown et al. 2010; Malbrán et al. 2014). During our observations, we noticed that the total number of spikelets varied among the spikes (data not shown). Therefore, the percent severity may vary slightly when the same number of diseased spikelets is observed in heads of different sizes. The advantages and technical (statistical) issues of using count or proportion data is a topic that deserves further investigation.

We provide data on the tebuconazole sensitivity for five FGSC-chemotype found in southern Brazil, but we acknowledge the limitations of our sample sizes. Thus far, few studies evaluated the overall fungicide sensitivity of at least two FGSC members from wheat in Brazil and used either small or more representative sampling (>50 strains) (Spolti et al. 2012a; Spolti et al. 2012b). The range of EC50 values found in our study varied not only among, but also within isolates of a same species, but they were within the range reported for *F. graminearum* isolates from the same region (Spolti et al. 2012a). Previously, differences in tebuconazole sensitivity between 3-ADON and 15-ADON strains within *F. graminearum* from NY State were not found when using large sample (25 strains each) (Spolti et al. 2014b). Although our small sample, which is far from being representative of the species, *F. graminearum* was in general less sensitive to tebuconazole because of one less sensitive isolate, than the other less frequent species. This agrees specifically with a recent study from our group where a larger sample of 15-ADON (48 strains) and the other two genotypes (NIV + 3-ADON = 17 strains) from barley was used (Machado et al. 2017). Conversely, in Uruguay, the NIV-producing strains within *F. asiaticum* and *F. cortaderiae* were significantly less sensitive than 15-ADON *F. graminearum* strains. Further molecular studies would be important to investigate the genetic basis of such differences in triazole sensitivity among the FGSC members.

Our results contribute new knowledge to improve our understanding of the ecology and epidemiology of *F. graminearum* species complex members, which may be of value for improving models for assessing the risk or epidemics and mycotoxin production. These should include assessing whether there are species-specific environmental requirements for fungal infection, colonization and mycotoxin production.

## Acknowledgements

First author is grateful to the Programa de Pós-graduação em Fitotecnia (UFRGS) and CNPq - Conselho Nacional de desenvolvimento Científico e Tecnológico, for providing a graduate scholarship CNPq.

---

**Figure 1S.**
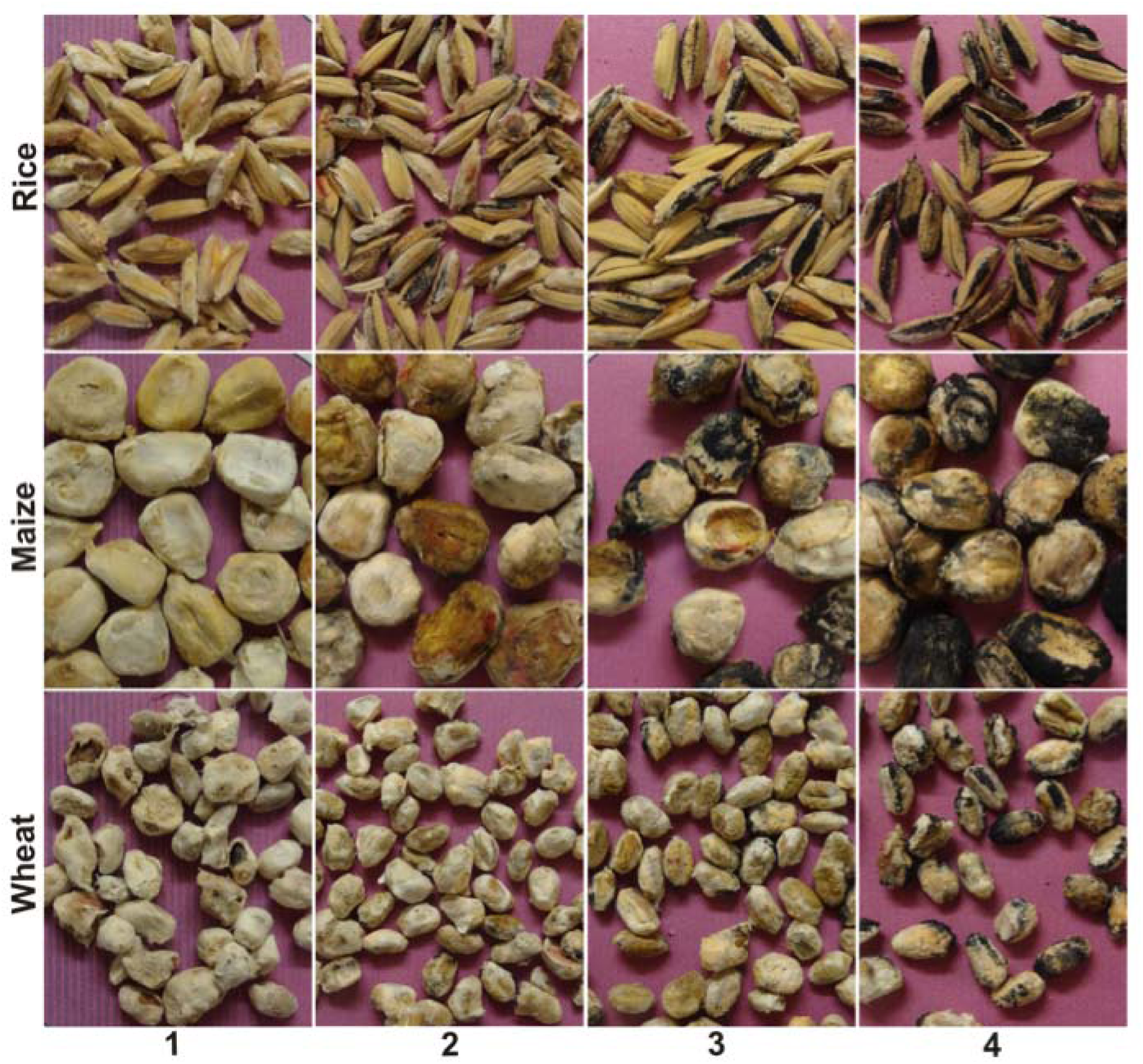
Diagrammatic key with four scores representing a percent range for the grain surface covered with perithecia, for rice, maize and wheat substrates. 1 = 0% (no perithecia); 2 = 1-10% ; 3 = > 10% - 30%); 4 = > 30%.

